# SARS-CoV-2 N-protein induces the formation of composite α-synuclein/N-protein fibrils that transform into a strain of α-synuclein fibrils

**DOI:** 10.1101/2023.03.13.532385

**Authors:** Slav A. Semerdzhiev, Ine Segers-Nolten, Paul van der Schoot, Christian Blum, Mireille M.A.E. Claessens

**Affiliations:** Nanobiophysics, Faculty of Science and Technology, MESA + Institute for Nanotechnology and Technical Medical Centre, University of Twente, P.O. Box 217, 7500 AE Enschede, The Netherlands; Theory of Polymers and Soft Matter, Eindhoven University of Technology, P.O. Box 513, 5600 MB Eindhoven, The Netherlands

## Abstract

The presence of deposits of alpha-synuclein fibrils in cells of the brain are a hallmark of several α-synucleinopathies, including Parkinson’s disease. As most disease cases are not familial, it is likely that external factors play a role in disease onset. One of the external factors that may influence disease onset are viral infections. It has recently been shown that in the presence of SARS-Cov-2 N-protein, αS fibril formation is faster and proceeds in an unusual two-step aggregation process. Here, we show that faster fibril formation is not due to a SARS-CoV-2 N-protein-catalysed formation of an aggregation-prone nucleus. Instead, aggregation starts with the formation of a population of mixed αS/N-protein fibrils with low affinity for αS. After the depletion of N-protein, fibril formation comes to a halt, until a slow transformation to fibrils with characteristics of pure αS fibril strains occurs. This transformation into a strain of αS fibrils subsequently results in a second phase of fibril growth until a new equilibrium is reached. Our findings point at the possible relevance of fibril strain transformation in the cell-to-cell spread of the αS pathology and disease onset.

## Introduction

The presence of deposits of alpha-synuclein fibrils in cells of the brain are a hallmark of α-synucleinopathies, including Parkinson’s disease, dementia with Lewy bodies and multiple system atrophy. Alpha-synuclein is a 140 amino acid intrinsically disordered protein. It is abundantly present in the brain, where it plays a role in membrane remodeling and membrane trafficking.^1–6^ For reasons that are not well understood, αS loses its physiological function and self-assembles into amyloid fibrils. These fibrils form deposits in disease specific brain cells and regions. It has been suggested that, similarly as in prion disease, the different pathologies are the result of αS amyloid fibrils with different structural characteristics, also referred to as *fibril strains*.^7,8^ The fibril strain is conserved in the cell-to-cell transmission of the amyloid fold and the progression of the disease.

The intrinsic architecture of the amyloid fold is replicated and preserved between cells.^7^ Which αS fibril strain is found in brain cells therefore likely depends on how the aggregation starts. Various experiments have shown that different external factors, including the presence of (poly)cations, virus proteins, lysozyme fibrils, and fine particulate matter affects αS aggregation.^9–13^ Since most cases of α-synucleinopathies are not familial, external factors and/or changes in internal factors due to aging likely play a dominant for disease onset.

One of the external factors that may influence disease onset is viral infection. For some viruses this is well established in case studies or on an epidemiological level.^14–18^ In the context of the covid pandemic, first cases that connect SARS-CoV-2 infections to the development of Parkinson’s disease (PD) have indeed been reported. We have recently presented results that point toward direct interactions between the N-protein of SARS-CoV-2 and α-synuclein as a molecular basis for the observed relation between SARS-CoV-2 infections and Parkinsonism. In the presence of N-protein αS fibril formation is not only much faster, it also proceeds in an unusual two-step aggregation process in which two different fibril strains are formed. Here we show that the observed aggregation kinetics result from the initial formation of a population of mixed αS/N-protein fibrils with low affinity for αS. After the depletion of N-protein, fibril formation grinds to a halt until a slow strain transformation to a pure αS fibril strain occurs. This pure αS fibril strain has a higher affinity for αS and dominates the fibril population at later times. Upon subsequent addition of monomeric αS the first strain only slowly elongates or requires the presence of additional N-protein. The second fibril strain readily seeds αS aggregation. In view of cell-to-cell spread of the αS pathology we therefore hypothesize that this strain conversion is required for transmission of the amyloid fold.

## Results and Discussion

Aggregation assays show that the aggregation of αS proceeds considerably faster in the presence of the SARS-CoV-2 N-protein. Moreover, a two-step aggregation process is observed (Fig 1a).^10^ In the presence of the amyloid reporter dye ThioflavinT (ThT), we observe an increase of ThT emission after a lag time, t0, to a first plateau, p1. After some period of time in which the ThT intensity remains constant, the ThT emission increases again to a final plateau, p2. The nature of the two plateaus and of the transition is unknown. To characterize the fibrils formed at the plateaus p1 and p2 and those during the transition from p1 to p2, we recorded AFM images at different times. From these AFM images we obtained the helical pitch via a discrete Fourier transform. We observe that going from p1 to p2 the helical pitch of the fibrils changes. Initially, in p1 fibrils with a helical pitch of approximately 86 nm dominate. In time, increasing number of fibrils with a pitch of 126 nm appear. Finally, in p2 fibrils with a pitch of approximately 126 nm dominate (Fig 1b, c). We refer to the fibrils with a pitch of 86 and 126 nm as different fibril strains, s1 and s2, respectively.

**Figure 1.**
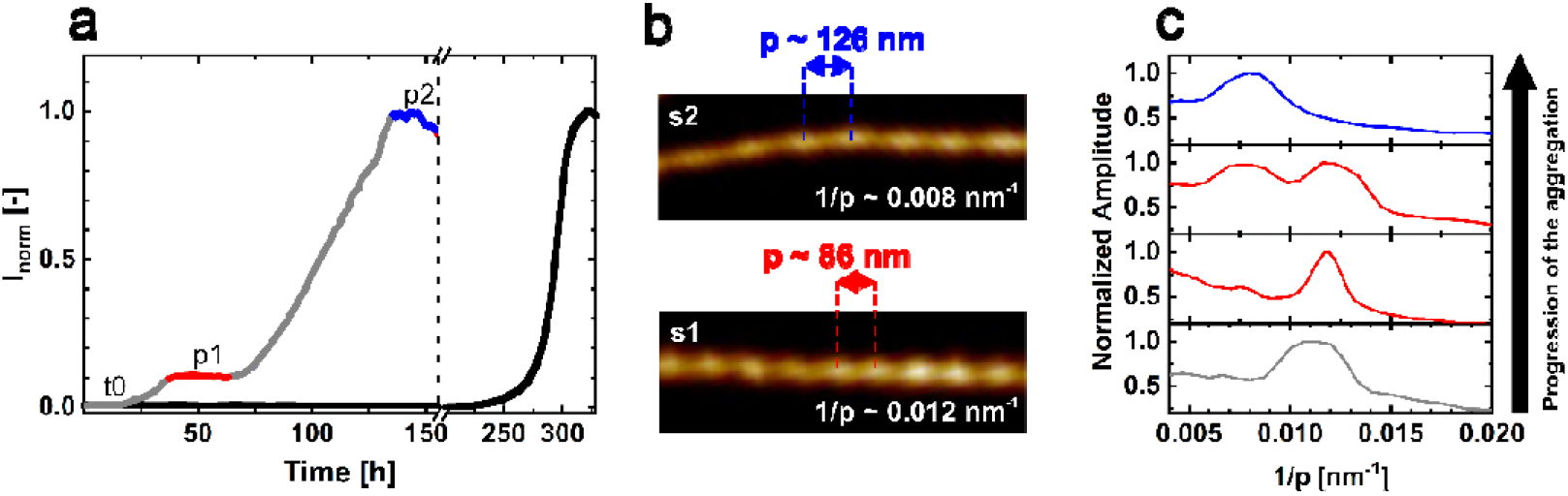
Aggregation of αS in the presence N-protein. **a)** Aggregation assay of αS in the absence (black) and presence (grey) of SARS-CoV-2 N-protein. The aggregation process is followed by recording the fluorescence of the amyloid-binding dye ThT. The aggregation of αS in the presence of N-protein is much faster compared to the aggregation in the absence of N-protein. In the presence of N-protein, aggregation proceeds in two steps. After a lag time, t0, a plateau, p1 (red) is reached, and aggregation resumes after some time lag to second plateau, p2, (blue). The dashed line marks the break in the x-axis. **b)** AFM images of fibrils obtained from the stages p1 and p2. In p1 and p2 two different fibril strains s1 and s2 are observed. The mean periodicity p of s1 is 86 nm and of s2 is 126 nm. **c)** Discrete Fourier analysis on fibrils obtained during the aggregation from p1 to p2. Initially, s1 fibrils with a periodicity of 86 nm dominate. In time, s2 fibrils appear, which is visible as a growing peak at 1/p=0.008 nm^−1^. In stage p2 the s2 fibril strain dominates.

**Figure 2.**
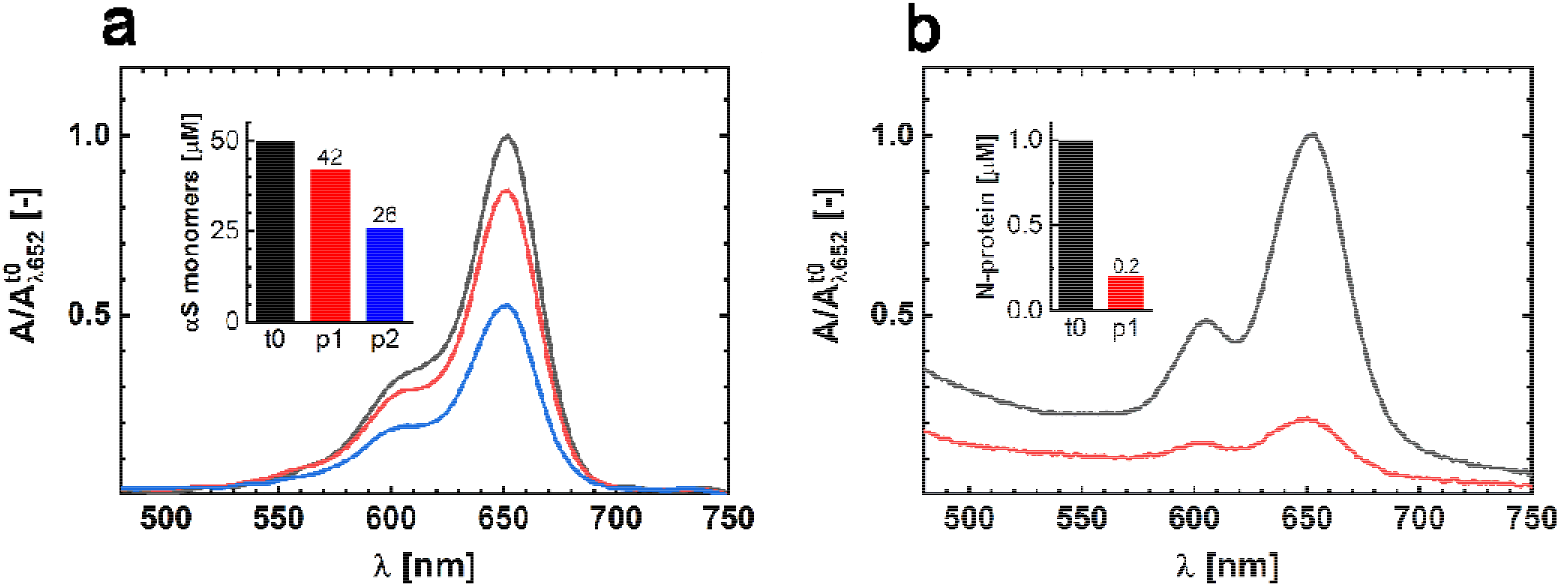
Residual αS and N-protein monomers concentration in solution. **a)** Absorbance spectra of αS-AF647 in the supernatant after spinning down the fibrils. Data are shown for samples obtained in the stages t0 (black), p1 (red) and p2 (blue). The data are peak-normalized to the t0-stage absorbance. The inset shows the derived total residual αS monomer concentrations at the different stages of the aggregation process. **b)** The experiment was repeated with N-protein-AT647. The absorbance spectra are recorded from the supernatant after spinning down the fibrils. Data are shown for samples obtained in the stages t0 (black) and p1 (red). The data are peak-normalized to t0 absorbance. The inset shows the derived total residual N-protein concentrations at the different stages of aggregation.

It is not clear what the origin of the two different ThT intensity plateaus is. It is well known that the fluorescence intensity of ThT bound to amyloid fibrils not only depends on the fibril mass but also on the fibril morphology.^19^ The two plateaus could thus just represent the two different fibril strains. To directly determine the fibril mass we measured the residual αS monomer concentration in solution. To be able to discriminate between αS and N-protein we used a fraction of fluorescently labelled αS in the aggregation. Samples were taken from the p1 and p2 plateaus, and the formed amyloid fibrils were spun down via high speed centrifugation. The residual monomer concentration in the supernatant was determined by measuring the absorbance of the fluorescently labelled αS-AF647 at 650 nm and using the Lambert-Beer relation. In stage p1, we find a residual monomer concentration of 42 μM, whilst in stage p2 we measure a concentration of 26 μM αS. In p1, 8 μM of αS has aggregated into fibrils while in p2 the amount of αS aggregated increases by a factor three, to 24 μM. The fibril mass clearly increases going from p1 to p2, showing that an increase in the total fibril mass accompanies the two-step aggregation. However, the increase in ThT intensity is not proportional to the increase in fibril mass, which also supports the finding that p1 and p2 contain different fibril strains. The residual αS monomer concentrations at p1 and p2 show that the affinity for αS monomer addition is different for the two strains. First, a strain with lower αS affinity is formed, subsequently a fibril species with higher αS affinity appears. The plateaus in the ThT intensity trace represent the apparent equilibrium between amyloid fibrils and monomers.

The N-protein induces the aggregation of αS. It has been shown that this proceeds via the formation of αS/N-protein complexes.^10^ If N-protein only acts as a catalyst that triggers aggregation, or plays a direct role in the formation of fibrils, remains unclear. To determine whether the N-protein is consumed in fibril formation, we use the same approach outlined above and determined the residual N-protein concentration in p1. In p1 we observe a strong depletion of N-protein in the supernatant. Absorbance measurements show that in p1 only 20% of the starting N-protein concentration remains in the supernatant (fig.1e) evidencing that most of the N-protein is consumed in the aggregation process. We therefore conclude that N-protein does not act as a catalyst, as the N-protein is not released after inducing αS aggregation.

In p1, αS and N-protein have been consumed in a ratio of approximately 10 to 1, and the supernatant is depleted of N-protein, while αS is still present in excess. If N-protein is a reaction partner in fibril formation, it should therefore be possible to restart the formation of s1 fibrils. The addition of additional N-protein in p1 indeed results in an restart of fibril formation as evidenced by an immediate increase in ThT intensity. The increase in the ThT signal levels off with time, as observed for the initial increase to p1. Fibril formation continues until N-protein is depleted from solution again. The newly formed fibril mass scales with the added amount of N-protein. Using fluorescently labelled N-protein in the aggregation confirms that N-protein is part of the formed fibrils. We observe a colocalization of fluorescence from the labelled N-protein and ThT (Fig. 3b). To exclude that N-protein forms amyloid fibrils by itself we isolated the s1 fibril to remove the residual αS and added N-protein. The isolated s1 fibrils do not seed the formation of pure N-protein fibrils (Fig. 3c). Combined these results show that both N-protein and αS are required for the formation of s1 fibrils. At p1, composite fibrils containing both αS and N-protein have been formed. The formation of composite fibrils containing N-protein is in line with reports on the formation of amyloid-like fibrils of N-protein in the presence of viral RNA. ^20^ It has been observed that interaction between the low-complexity domain of the N-protein and viral RNA facilities fibril formation. The net negatively charged disordered αS may act in a similar way, resulting in composite αS/N-protein fibrils.

**Figure 3.**
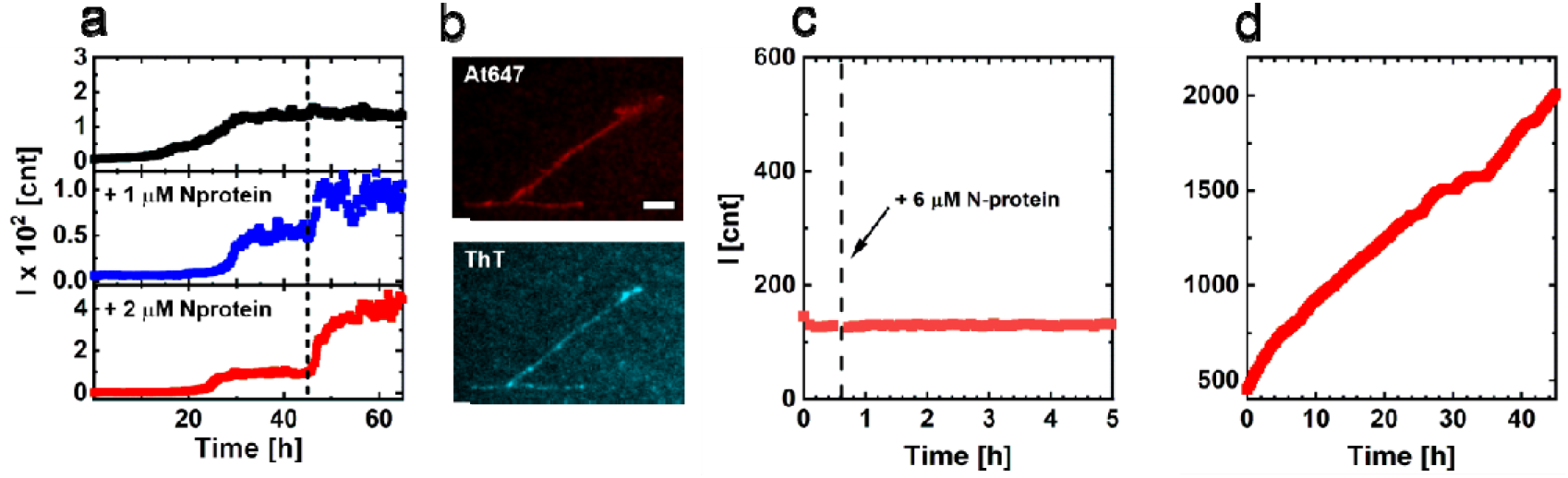
**a)** After reaching the plateau p1, additional N-protein was added at the time point indicated by the dashed line, no N-protein added in black, 1 μM N-protein added in blue, 2 μM N-protein added in red. The addition of N-protein immediately restarts the aggregation. The increase in ThT emission reflects the added amount of N-protein. **b)** Fluorescence microscopy image of fibrils formed in the presence of At647 labelled N-protein. The top panel show the emission from the At647 labelled N-protein. The bottom panel shows the ThT emission and confirms that the visualized structures are amyloid fibrils. **c)** ThT assay of resuspended isolated s1 fibrils. At the timepoint indicated by the dashed line N-protein was added. The addition of N-protein did not restart aggregation. **d)** ThT assay of αS monomers in the presence of resuspended isolated s2 fibrils under quiescent conditions. Addition of the s2 fibrils results in a linear increase of the ThT emission with time evidencing that the increase in fibril mass is dominated by fibril elongation.

Since N-protein has been depleted in p1, while αS was still present in excess, the s2 fibrils must predominantly contain αS. Indeed, the characteristics of the s2 fibrils resemble the characteristics of αS fibrils formed in the absence of N-protein under comparable conditions. The fibril pitch and the residual monomer concentration matches values found for pure αS fibrils.^21,22^ To test if the s2 can seed αS aggregation we added 50 μM monomeric αS and let the aggregation proceed under quiescent conditions. The addition of αS to s2 fibrils results in an immediate start and linear increase of the ThT intensity with time (Fig. 3d). Clearly the s2 fibrils seed αS aggregation. The linear increase in ThT intensity with time evidences that the increase in fibril mass is dominated by the addition of αS monomers as expected under these conditions.

The depletion of N-protein is the reason for the observed plateau p1. Next, we investigated the mechanism that underlies the restart of the aggregation and the formation of s2 fibrils. The formation of two different fibril strains in a two step aggregation process has been observed before. Aggregation of a mixtures of the amyloid peptides Aβ40 and Aβ42 results in an apparent two-step aggregation behavior due to the formation of two different fibril strains, which independently form following different lag times.^23^ However, in our experiments the aggregation towards p2 is fast compared to de novo formation of αS fibrils (Fig. 1a). We therefore rule out that the two-step aggregation process is the result of two independent nucleation processes of composite αS /N-protein fibrils and pure αS fibrils. Another mechanism that could account for the observed two-step aggregation process is surface catalyzed secondary nucleation (SCSN) of amyloids, where the surface of the composite fibrils nucleate the formation of pure αS fibrils.^24,25^ To test if SCSN is responsible for the observed behavior, we performed the aggregation of αS at quiescent conditions in the presence of s1 fibrils. The experiments were performed in excess of αS at different αS concentrations. At the start of the experiment we observe a linear increase of ThT signal for all αS concentrations tested (Fig. 4). Upon depletion of αS, the curves level off, and the ThT intensity scales with the αS concentration added. The plateau ThT intensity hence reflects the total mass of the fibrils. The initial linear increase in ThT intensity with time points at a growth of the fibrils from the fibril ends. If the aggregation kinetics is governed by the rate of addition of monomers to the fibril ends, the aggregation can be treated as a bimolecular reaction of first order.^22^ When we plot the initial slope of the ThT curves as a function of the added αS monomer concentration we indeed observe a linear relation (Fig. 4b). This linear increase of the initial growth rate with the added αS monomer concentration signifies that addition of αS to s1 fibrils results in a bimolecular reaction of the first order. We therefore conclude that the transition into the second plateau is *not* caused by the nucleation of new fibrils via SCSN.

**Figure 4.**
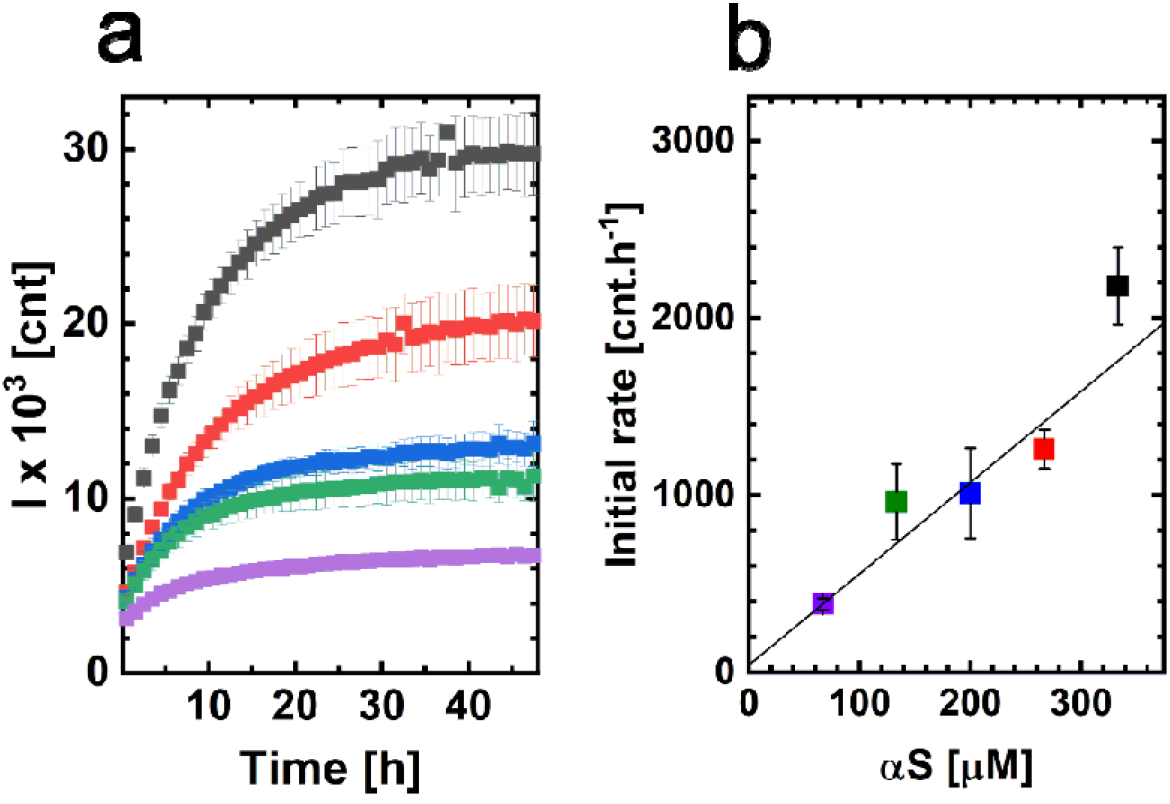
Aggregation of αS in the presence fibrils from p1. **a)** Increase in ThT emission as a function of time for increasing concentration of added monomeric αS, 333 μM (black), 267 μM (red), 200 μM (blue), 133 μM (green), 67 μM (purple) and constant concentration (~4 μM equivalent monomer) of p1 fibrils. **b)** Initial slope of the ThT emission increase observed for the different added αS monomer concentrations. The color coding is the same as in a).

In the absence of new nucleation events, the fibrils themselves have to transform, and the obtained data suggests that s1 fibrils elongate and transform to the s2 phenotype. To directly visualize this fibril growth, we started the aggregation of αS in the presence of N-protein with a fraction of αS labelled with αS-AF647 under mild agitation. After reaching p1, αS conjugated with αS-AF568 (Fig. 3 schematic) was added and aggregation was allowed to proceed. Fluorescence microscopy images of the resulting s2 fibrils are shown in Figure 5. The images reveal the presence of segments of different color, which confirms that the fibrils that are formed in p1 elongated upon transition to p2. Besides fibrils that contain segments of different color we also observe fibrils of only one color. We attribute the presence of one color fibrils to fibril breaking. We conclude that the restart to the second plateau is caused by a strain transformation towards the s2 morphology within the elongating fibrils. This strain transformation is a chance process, the increase in fibril mass is initially very slow, resulting a near constant ThT signal in p1. With the occurrence of a strain transformation the fibrils elongate up to a point where sample agitation causes the elongated fibrils to break, creating more fibrils. The increase in the number of ends results in an exponential increase of the fibril mass towards p2.

**Figure 5.**
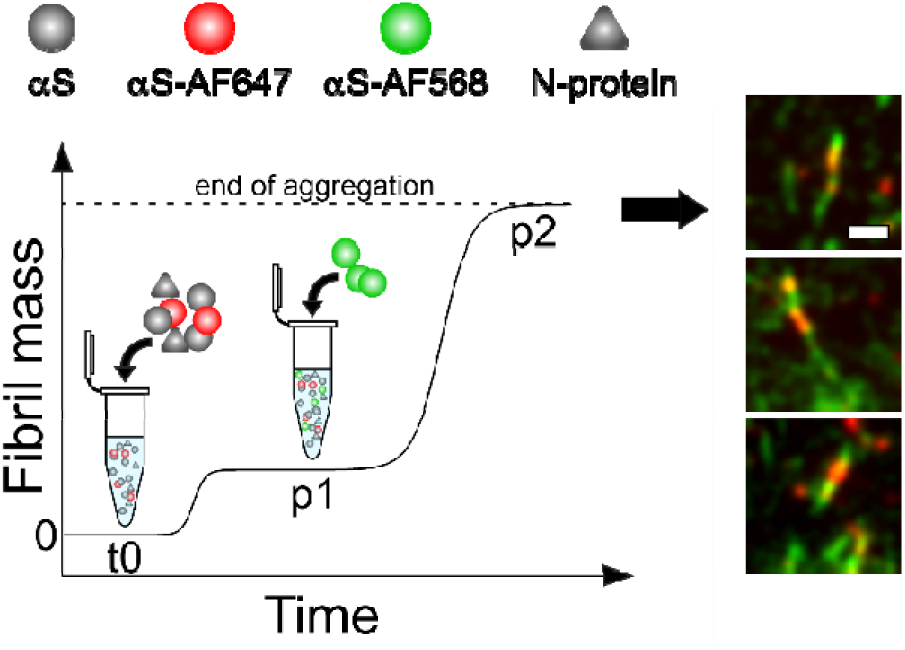
Visualizing the two steps of αS aggregation in the presence of N-protein. The starting αS/N-protein aggregation mixture in stage t0 contains a fraction of αS-AF647 (red). Once the first plateau p1 is reached, αS-AF568 is added (green). The resulting fibrils were imaged using TIRF microscopy after p2 was reached. The resulting fibrils clearly show segments of different color. This evidences that the s1 fibril strains extends and converts into the s2 fibril strain. Scale bar is 2 μm.

## Conclusions

In summary, the SARS-CoV-2 N-protein does not induce αS aggregation by catalysing the formation of an aggregation-prone nucleus. Instead, N-protein and αS co-aggregate resulting in the formation of a composite αS/N-protein fibril. These composite fibrils have relatively low affinity for αS and do therefore not deplete the solution of αS. After reaching the equilibrium between αS, N-protein and the composite fibril, the increase in fibril mass comes to an apparent stop. However, at the fibril ends a slow strain conversion occurs upon the addition of αS monomers. This transformation into a strain of αS fibrils subsequently results in a second phase of fibril growth until a new equilibrium is reached. The fibrils of the second strain resemble pure αS fibrils and have a much higher affinity for the αS monomers. In view of the observed relation between different αS pathologies and αS fibril both the fast formation of the first composite fibril strain, as well as the subsequent strain transformation to a αS fibril strain, may be of interest. The composite fibril strain requires the presence of N-protein to propagate and can therefore not directly induce fibril formation in cells that are not infected with SARS-CoV-2 and that therefore do not express N-protein. However, after the strain transformation to the αS strain, N-protein is no longer required for propagation. This new strain can therefore also induce fibril formation in cells that are not infected with SARS-CoV-2. Note, that formation of the αS strain from the composite strain is much faster than *de novo* formation of αS fibril. Our data point at the possible relevance of fibril strain transformation in the cell-to-cell spread of the αS pathology and disease onset.

## Materials and methods

### Preparation and Labelling of αS Monomers

Expression of the human αS wild-type (αS), the αSA140C and αSA18C mutants with a single alanine to cysteine substitution, was performed in E. coli B121 (DE3) using the pT7-7-based expression system. Details on the purification procedure for αS and RS140C are described elsewhere. ^26^ The αSA140C and αSA18C mutants were conjugated with AlexaFluor 647 and AlexaFluor 568 NHS-esters (Thermo Fisher Scientific, USA) targeting the cysteine residues. Both αS variants were first incubated with 2 mM DTT to ensure the reduction of any present disulfide bridges. The DTT was removed using a 2 ml ZebaSpin MWCO 7 kD desalting column (Thermo Fisher Scientific, USA). The columns were pre-equilibrated by washing them with 1 ml 10 mM TRIS buffer at pH 7.5 for three consecutive times by spinning them at 1000 G for 2 minutes for each washing step. The protein solution was placed in the equilibrated column and was spun down for 2 minutes at 1000 G. The DTT-free eluate containing the αS variant was collected. 100 μM of each αS variants was incubated with 2-fold excess of the respective AlexaFluor dye at room temperature, in dark and under mild agitation for the duration of two hours. The reaction mixture was then transferred to 2 ml ZebaSpin MWCO 7 kD (Thermo Fisher Scientific, USA) columns to separate the labeled protein from the free dye. The columns were washed 3 times with 10 mM TRIS buffer at pH 7.5. The reaction mixture was spun down for 2 minutes at 1000 G and the eluate containing the labeled protein was collected for further characterization. The degree of labeling (DOL) was determined by means of UV-Vis absorbance spectroscopy. The DOL was 0.5 and 1 for αS-AF647 and αS-AF568 respectively.

### Preparation and labelling of SARS-CoV-2 N-protein

The SARS-CoV-2 N-protein was recombinantly produced in our laboratory following the procedure mentioned elsewhere.^27^ The N-protein was labeled using Atto 647 NHS - ester (Atto-Tec, Germany) targeting the accessible amino groups of the protein. 40 μM N-protein in 20 mM Hepes, pH = 8, 100 mM NaCl was incubated with 3-fold excess of Atto 647 NHS at room temperature, under gentle agitation, for 2 hours in the dark. The reaction mixture was then transferred to 2 ml ZebaSpin MWCO 7 kD desalting column (Thermo Fisher Scientific, USA) to separate the labeled protein from the free dye. The columns were pre-equilibrated by washing them with 20 mM Hepes, pH = 8, 100 mM for three consecutive times by spinning them at 1000 G for 2 minutes for each washing step. The reaction mixture was spun down for 2 minutes at 1000 G and the eluate containing the labeled protein was collected for further characterization. The degree of labeling (DOL) of ~1. 8 was determined by means of UV-Vis absorbance spectroscopy.

### THT Aggregation Assays

In the aggregation assays, a solution of 50 μM αS and 1 μM N-protein in 20 mM Tris buffer (Sigma-Aldrich, UK), pH = 7.4, 5 μM ThT (Fluka, Sigma-Aldrich, UK), and 10 mM NaCl (Sigma-Aldrich, USA) was placed in wells of a 96-well half area clear flat-bottom polystyrene NBS (low bind) microplate (3881, Corning, US). Samples were prepared in 3 replicates of 50 μL. The aggregation proceed while shaking at 500 rpm at 37 °C. To follow the aggregation, the increase in ThT fluorescence was monitored using a plate reader (Infinite 200 Pro, Tecan Ltd., Switzerland). The ThT dye was excited at 446 nm and the fluorescence signal was measured at 485 nm every 10 min.

### Atomic Force Microscopy

The samples from the aggregation mixture (50 μM αS, 1 μM N-protein) were diluted five times, deposited onto mica (Muscovite mica, V-1quality, EMS, US), and left to rest for 5 minutes. The sample was carefully washed 3 times with 20 μL of demineralized water (Milli-Q) and gently dried under flow (low flow rate) of nitrogen gas. AFM images were obtained using a BioScope Catalyst (Bruker, US) in soft tapping mode using a silicon probe, NSC36 tip B with a force constant of 1.75 N/m (MikroMasch, Bulgaria). Images were captured with a resolution of 512 × 512 (10 μm × 10 μm) pixels per image at a scan rate of 0.2 to 0.5 Hz. AFM images were processed with the Scanning Probe Image Processor (SPIP, Image Metrology, Denmark) and the Nanoscope Analysis (Bruker, US) packages. The fibril morphology was analyzed using a custom fibril analysis Matlab script adapted from the FiberApp package.^28^

### Determination of the residual monomer concentration

The residual αS monomer concentration was determined by first aggregating 49 μM αS, 1 μM αS-AF647 (DOL = 0.5) and 1 μM in conditions identical to those described in the Aggregation assays section. Samples where taken out of the well plate at either p1 or p2 and were spun down for 1 hour at 18 kG. The absorbance of the supernatant was measured at λ = 650 nm using the Nanodrop ND-1000 (ThermoFisher Scientific, USA) or the UV −2401 (Shimadzu, Japan) UV-Vis spectrophotometer. The concentration of αS-AF674 was determined spectroscopically and from this the total residual αS monomer concentration was derived.

The residual N-protein concentration was determined by first aggregating 50 μM αS, 0.5 μM N-protein and 0.5 μM N-protein-At647 (DOL = 1.8) at conditions identical to those described in the Aggregation assays section. Samples where taken out of the well plate at either p1 or p2 and incubated at high-ionic strength conditions (0.5 M NaCl) for 1 hour to minimize adsorbance of the net positively charged N-protein to the fibrils which are mainly composed of net negatively charged αS. The remainder of steps to determine residual N-protein concentrations are identical to those taken for αS (see above).

### Aggregation in the presence of fibrils from p1 and p2

#### αS and N-protein monomers in the presence of fibrils from p1

αS and N-protein were aggregated following the protocol described in the Aggregation assays section. After p1 was reached the N-protein concentration was increased by ~1 μM and ~2 μM from a 40 μM N-protein stock solution in 20 mM Tris buffer. After the addition of N-protein monomers the aggregation assay and the readout commenced in the plate reader.

#### N-protein monomers in the presence of fibrils from p1

αS and N-protein were aggregated following the protocol described in the Aggregation assays section. When p1 was reached, the aggregation mixture was taken out of the well plate and spun down for 1 hour at 18 kG. The pellet with p1 fibrils was resuspended in 20 mM Tris buffer, 10 mM NaCl. The solution of resuspended p1 fibrils was placed in a well and the well plate was returned to the plate reader were the aggregation was reinitiated at the same conditions as the ones in which the p1 fibrils were produced. After ~45 min the N-protein concentration was increased by 6 μM. After the addition of extra N-protein the aggregation and readout was continued.

#### αS in the presence of fibrils from p1

A solution of 50 μM αS, 1 μM N-protein, 20 mM Tris, pH = 7.5, 5 mM ThT, 10 mM NaCl was incubated in Thermomixer comfort (Eppendorf, Germany) at 37 °C and shaking at 500 RPM. The aggregation was followed by measuring the ThT emission signal at λ = 485 nm using a Cary Eclipse Fluorescence Spectrometer (Agilent Technologies, USA). When p1 was reached, the aggregation mixture containing the p1 fibrils was used to prepare the samples for the aggregations. In the final samples, the solution obtained in p1 was diluted 3 times and varying αS monomer concentrations were added to reach the set of concentrations that given in the main text. Besides the αS concentration experimental conditions remained identical to one used for the initial aggregations. The samples were placed in a well plate. The aggregation and the read out were performed on Saffire^2^ (Tecan, Switzerland) under quiescent conditions, at 37 °C, excitation at 445 nm and emission at 485 nm. The ThT signal was recorded every 15 min. The initial rates were determined using the data points recorded within the first 2 h of the aggregation.

#### αS monomers in the presence of fibrils from p2

αS and N-protein were aggregated following the protocol described in the Aggregation assays section. When p2 was reached, the aggregation mixture was taken out of the well plate and spun down for 1 hour at 18 kG. The pellet of p2 fibrils was resuspended in 20 mM Tris, 10 mM NaCl and monomeric αS was added to increase the total concentration with 50 μM. The solution then was placed back in well and the well plate returned to the plate reader. The aggregation and read out were reinitiated at quiescent conditions at 37°C

### Microscopy

#### Visualizing the two steps of αS aggregation in the presence of N-protein

50 μM αS, 0.14 μM αS-AF647 and N-protein were aggregated following the protocol described in the Aggregation assays section. When p1 was reached the aggregation was paused 0.16 mM of αS–AF568 was added. Then the aggregation was continued at the same conditions. Once the aggregation was finished (p2) the resulting fibrils were imaged using Nikon TiE in total internal reflection fluorescence (TIRF) mode. For the excitation of the two fluorescent labels the 514 nm line of an argon laser (35-IMA-040, Melles Griot, USA).) and a 640 nm diode pumped solid state laser (Coherent Cube 640 100C, US) were used. To image αS-AF568, a filter cube containing a 514 nm excitation filter with 10 nm bandpass, a 514 nm dichroic and a 525 long pass emission filter was used. To image αS-AF64, a filter cube containing a 640 nm excitation filter with 20 nm bandpass, a 640 nm dichroic and a 675 nm emission filter with 50 nm bandpass was used. The images were obtained using a lOOx Apo TIRF 1.49 oil objective (Nikon, Japan) and recorded with an iXon 3 Andor DU-897 EMCCD camera (Oxford instruments, UK).

#### Imaging ThT stained fibrils containing N-protein-At647

50 μM αS and 1 μM N-protein-At647 were aggregated following the protocol described in the Aggregation assays section. Once the aggregation reached p1 the resulting fibrils were imaged using Nikon TiE TIRF microscope with identical optical setup to the one used above (see Visualizing the two steps of αS aggregation section) To excite the ThT dye, the 457 nm line of an Argon laser (35-IMA-040, Melles Griot, USA) was used. The filter cubes used for ThT contained a 455 nm excitation filter with 10 nm bandpass, a 458 nm long pass dichroic mirror and a 485 nm emission filter with 30 nm bandpass. The filter cube used for αS-AF647 was used also for N-protein-At647.

## The Acknowledgments

We would like to thank Kirsten A. van Leijenhorst-Groener for the production of the recombinant proteins. We are grateful to the Dutch Parkinson’s disease foundation “Stichting ParkinsonFonds” for financial support.

